# Altiratinib Targets PRP4K in *Theileria annulata*, Disrupting RNA Processing and Inducing Apoptosis in Infected Cells

**DOI:** 10.1101/2025.10.24.684341

**Authors:** Madhusmita Subudhi, Akash Suresh, Siva Singothu, Vasundhra bhandari, Sakshi Singh, A Vengatachala moorthy, Paresh sharma

## Abstract

Tropical theileriosis, driven by *Theileria annulata*, poses a critical threat to livestock health, particularly in the face of rising resistance to buparvaquone, the primary and only treatment option. This study investigates the therapeutic potential and mechanistic actions of Altiratinib, a spliceosome-associated kinase inhibitor, in targeting *T*.*annulata*-infected bovine cells. Through bioinformatic and molecular docking analyses, Altiratinib was shown to selectively bind conserved catalytic motifs of PRP4K homologs in *T. annulata* (TA21325) and *Theileria parva* (TpMuguga_01g00303), with reduced affinity observed upon L715F mutation. *In vitro* assays confirmed potent anti-parasitic activity against both buparvaquone-sensitive and - resistant strains, while sparing uninfected peripheral blood mononuclear cells. Proteomic profiling revealed disruption of host and parasite RNA splicing and translation machinery. Altiratinib further induced G1-phase arrest, inhibited cMET-mediated signaling, and suppressed DNA synthesis. Additionally, it triggered oxidative stress, mitochondrial depolarization, and ROS-mediated DNA damage, culminating in p53 activation and caspase-9-driven intrinsic apoptosis. Downregulation of TaSP expression reinforced its selective targeting of parasite-infected cells. These findings highlight Altiratinib’s promise as a directed therapeutic for tropical theileriosis and other piroplasm infections, offering a novel avenue to combat drug-resistant strains and meriting further preclinical evaluation.

**Graphical abstract:** 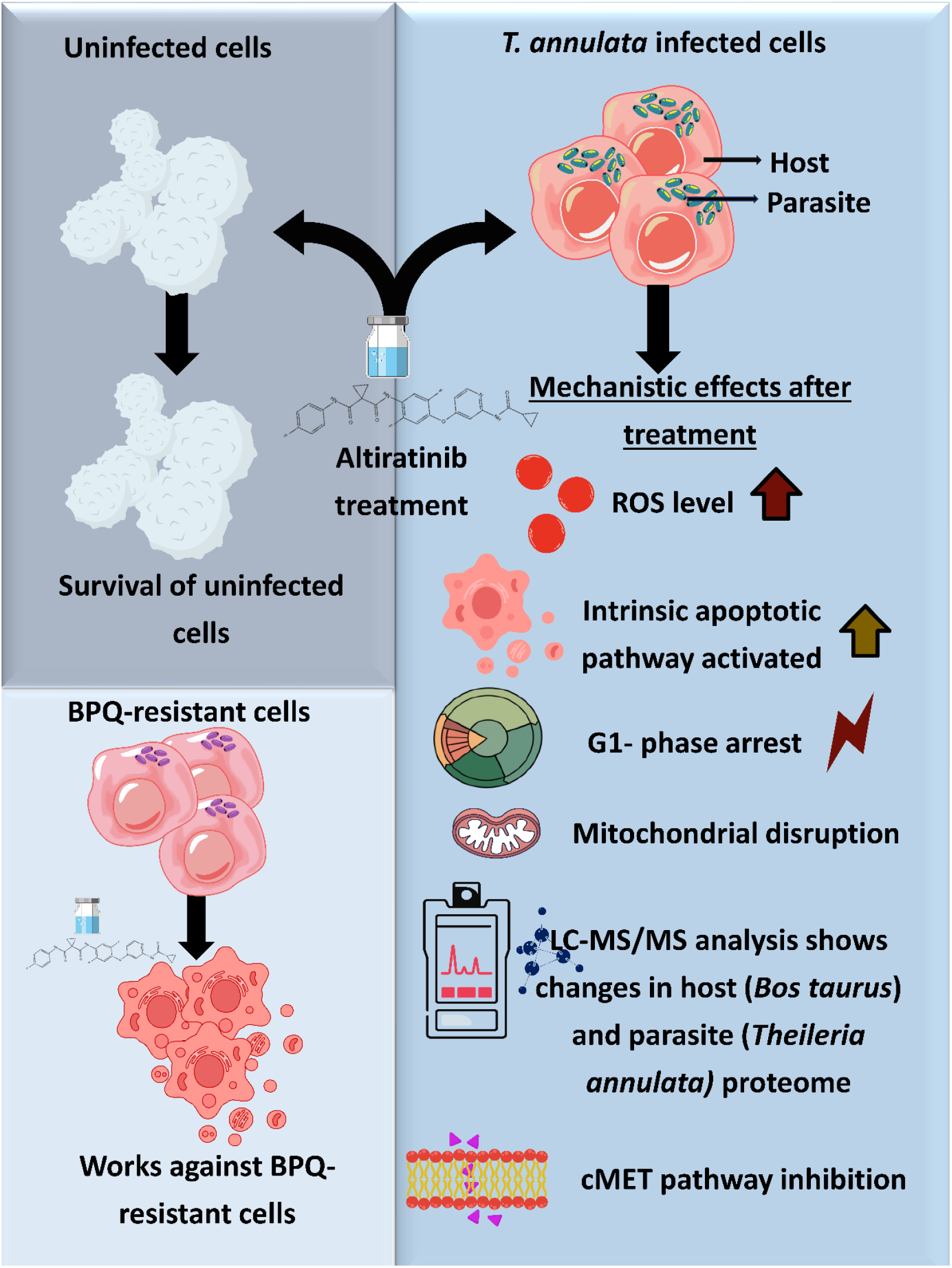

**Highlights:**

1. **Altiratinib disrupts spliceosomal and translational machinery** in *Theileria annulata*-infected host cells by targeting PRP4K homologs, leading to impaired RNA processing, oxidative stress, and intrinsic apoptosis.
2. **Demonstrates potent efficacy against both drug-sensitive and Buparvaquone-resistant strains**, validating its potential as a novel therapeutic strategy for overcoming resistance in tropical theileriosis.
3. **Dual-target mechanism** modulates parasite and host oncogenic pathways, including cMET signaling, revealing Altiratinib’s repurposing potential as a **broad-spectrum antiparasitic** with anticancer-like activity.

## 1. Introduction

The phylum Apicomplexa comprises obligate intracellular parasitic protozoa, including *Plasmodium, Toxoplasma, Theileria*, and *Babesia*, which are etiological agents of significant morbidity and mortality in both human and veterinary populations [1]. Infections caused by Apicomplexan parasites represent a significant global health challenge, contributing to high morbidity, mortality, and economic losses, particularly in resource-limited regions. Despite ongoing efforts to advance treatment options, therapeutic solutions for many Apicomplexa-related diseases remain limited, with some infections lacking effective treatments altogether [2]. Among these pathogens, *Theileria* and *Babesia* species are particularly notable for their role in tick-borne diseases that profoundly impact livestock health. These infections are endemic in regions such as sub-Saharan Africa, Asia, the Middle East, Southern Europe, and North Africa, where they pose a substantial threat to agricultural productivity and food security. *T. annulata*, the causative agent of tropical theileriosis, presents a major threat to cattle in countries such as India [3]. While *Buparvaquone* (BPQ) is the only treatment for *T. annulata* infections, the emergence of drug-resistant strains in regions like Tunisia, Iran, Turkey, India, and Sudan highlights the urgent need for new therapeutic options that target *T. annulata* through mechanisms less prone to resistance [4]. A key feature of *T. annulata* infection is its ability to manipulate host cell processes, inducing a cancer-like phenotype in macrophages and lymphocytes [5]. Despite progress in understanding these mechanisms, many aspects remain poorly understood, offering a potential avenue for developing novel therapeutic strategies. This study focuses on Altiratinib, a small-molecule inhibitor originally developed for cancer therapy and currently in phase II clinical trials (NCT02228811). Altiratinib targets critical signalling pathways, including c-Met (HGFR), VEGFR2, TIE2, and TRK, which play pivotal roles in tumor progression [6]. By inhibiting these pathways, Altiratinib effectively suppresses tumor cell proliferation and triggers apoptosis [7]. Recent research has highlighted its efficacy against *T. gondii* and *P. falciparum*, indicating its potential as a broad-spectrum therapeutic for Apicomplexan infections [8]. Given the evolutionary connections between *T. annulata, T. gondii*, and *P. falciparum*, Altiratinib emerges as a promising candidate for repurposing in treating *T. annulata* infections. This study explores the antiparasitic activity and mechanistic effects of Altiratinib in *T. annulata*-infected bovine cells, focusing on its dual impact on both the host and the parasite. Our findings reveal that Altiratinib disrupts the splicing machinery in the host and parasite, leading to oxidative stress, mitochondrial dysfunction, and cell cycle impairment. These results underscore Altiratinib’s potential as a novel therapeutic strategy for combating *Theileria* and other piroplasm parasites.

## 2. MATERIALS AND METHODS

### 2.1 Bioinformatics Analysis and Molecular Docking Simulations

A homology model of TA21325 was constructed using the protein sequence from the UniProt database (UniProt ID: Q4UGP2) in Schrodinger 13.1. To identify a suitable protein template, a BLASTp search was performed, revealing a 65% sequence identity with PDB ID: 7Q4A (chain A). The protein model was subsequently prepared using the Protein Preparation Wizard module of Schrodinger 13.1, where disulfide bonds and hydrogen bonds were formed, missing loops were filled, and water molecules were removed. Protein optimization and energy minimization were carried out using the OPLS4 force field. The receptor grid was generated to define the binding pocket using the Receptor Grid Generation module, Schrodinger 13.1. Altiratinib (PubChem CID: 54576299) was prepared by the LigPrep module in Schrodinger, converting the 2D structure into a 3D structure with generated tautomers at pH 7.0. The molecular docking between Altiratinib and the prepared protein was performed in extra precision mode to determine the docking score (kcal/mol). Binding affinity between Altiratinib and the TA protein was further analyzed using MM-GBSA (Molecular Mechanics/Poisson– Boltzmann Surface Area) in Schrodinger 13.1.

Conserved family domains were compared across different eukaryotes using the NCBI Conserved Domain Database (https://www.ncbi.nlm.nih.gov/Structure/cdd/wrpsb.cgi). Multiple sequence alignment and phylogenetic reconstruction were performed using ETE3 3.1.2 [9] through GenomeNet (https://www.genome.jp/tools/ete/). The phylogenetic tree was generated using FastTree with slow NNI and MLACC=3 [10]. Multiple sequence alignment was done using CLUSTAL W, and analysis was conducted using ESPript3.

### 2.2 Isolation, Culture of *T. annulata*-Infected PBMCs and *In Vitro* Evaluation of Drug Sensitivity

Peripheral blood samples were collected from cattle naturally infected with *Theileria annulata*. Peripheral Blood Mononuclear Cells (PBMCs) were isolated using density gradient centrifugation from whole blood and subsequently washed twice with phosphate-buffered saline (PBS). The isolated cells were cultured in complete RPMI-1640 medium (Life Technologies), supplemented with 10% fetal bovine serum (FBS) and 0.1% Penicillin-Streptomycin, under standard conditions (37 °C, 5% CO_2_). The *T. annulata*-infected lymphocytes (TA cells) exhibited continuous proliferation in vitro, indicative of parasite-mediated cellular immortalization. The establishment of *T. annulata*-infected cell lines was confirmed by PCR amplification targeting the *T. annulata* surface protein (TaSP) gene, following a few passages in culture, as previously described [11].

PBMCs were isolated from apparently healthy, *T. annulata*-negative cattle (n = 2) using the same protocol and were used as uninfected controls in subsequent assays. To evaluate the cytotoxic effect of Altiratinib on TA cell viability, dose-and time-dependent drug inhibition assays were performed. TA cells were seeded in 96-well plates at a density of 10,000 cells per well and treated with Altiratinib across a concentration range of 0.09–50 μg/mL. After 48 hours of drug exposure, cell viability was assessed using the resazurin reduction assay. Cells were incubated with 1.5 mM resazurin solution for 7–8 hours, and fluorescence was measured at an excitation wavelength of 570 nm.

To test the efficacy of Altiratinib against drug-resistant parasites, a buparvaquone (BPQ)-resistant TA cell line (BPQ-R) was developed. Parental TA cells, initially sensitive to BPQ at 200 ng/mL, were gradually exposed to increasing BPQ concentrations starting from 10 ng/mL. Through serial passaging, a resistant cell population capable of proliferating in up to 10 μg/mL BPQ was established. Notably, the drug-resistant phenotype was stable, as the cells maintained their resistance even after several weeks of culture in drug-free medium, without reverting to a sensitive state. BPQ was purchased from Sigma-Aldrich (B4725), while Altiratinib was obtained from Cayman Chemicals. Compounds were prepared as 1 mg/mL stock solutions in 100% DMSO.

### 2.3 Cell Proliferation Assay

A time-course study was conducted to assess the effects of Altiratinib on the proliferation of *Theileria*-infected cells. Cells (1 × 10^6^ per well) were treated with the IC_50_ concentration of Altiratinib and monitored for 48 hours. Cell counts were obtained every 12 hours using the trypan blue exclusion assay. The experiment was repeated five times with three replicates, and the average results were graphed.

### 2.4 Click Chemistry and EdU Incorporation Assay

The proliferation rate was evaluated using a Click-IT EdU Alexa Fluor 488 Imaging Kit (Thermo Scientific, #C10337). Cells were incubated with the IC_50_ concentration of Altiratinib for 48 hours and then treated with EdU for 3 hours. Following treatment, cells were fixed, permeabilized, and incubated with a Click-iT reaction cocktail. DNA was counterstained with Hoechst, and images were captured using a Zeiss fluorescent microscope with a 63× objective. Image processing and quantification were performed using ZEN 3.3 (blue edition) software.

### 2.5 Study of Cell Death Mechanisms and Apoptosis

To investigate cell death, *Theileria*-infected cells were treated with the IC_50_ concentration of Altiratinib for 48 hours. Apoptosis and necrosis were measured using Annexin V and propidium iodide (PI) staining. The experiment was conducted in duplicate to ensure reproducibility.

### 2.6 Analysis of Cell Cycle Phase Distribution and Mitochondrial Membrane Potential

For cell cycle analysis, cells (1 × 10^4^/mL) were treated with Altiratinib and stained with PI. After washing in PBS and fixation with ice-cold 70% ethanol, cells were incubated with cell staining buffer (50 μg/mL PI, 0.1% Triton-X, and 200 μg/mL RNase) at 37°C for 20 minutes. Cell cycle distribution was analyzed using BD flow cytometry and FlowJo X software. Mitochondrial membrane potential was assessed by staining cells with the JC-1 probe and comparing red-to-green fluorescence intensities using flow cytometry (BD LSR Fortessa) and FlowJo software.

### 2.7 Assessment of ROS Levels

Reactive oxygen species (ROS) levels were measured using H2DCFDA (Invitrogen #D399). After treatment with Altiratinib, cells were collected at various time points to analyze ROS production in a dose-dependent manner. Flow cytometric analysis was performed using a BD flow cytometer, and data were analyzed using FlowJo software.

### 2.8 RNA Isolation and Gene Expression Analysis

Total RNA was isolated using the Nucleospin RNA Extraction Kit (Macherey-Nagel). Reverse transcription to cDNA was performed with the Primescript cDNA Synthesis Kit (Takara, #6110A). Quantitative real-time PCR (qPCR) was carried out using SYBR Green Master Mix (Takara) to analyze relative gene expression. Gene expression levels were normalized to GAPDH, and primer sequences are provided in the Supplementary Table 1.

### 2.9 Immunofluorescence Analysis

Immunofluorescence assays (IFAs) were performed to assess oxidative damage (8-OHdG), phosphorylation of H2AX (Ser 139), p53 localization, and activation of caspases 8 and 9. Primary antibodies included Phospho-HistoneH2A.X (Ser139) (1:300), Phospho-p53 (Ser15) (1:500), Cleaved Caspase-9 (1:800), and Cleaved Caspase-8 (1:400), purchased from CST. After treatment with IC_50_ concentrations of Altiratinib, cells were fixed, permeabilized, and stained with primary and secondary antibodies. DNA was labelled with DAPI, and images were captured using a Zeiss fluorescent microscope. ZEN 3.3 (blue edition) software was used for image processing.

### 2.10 Western Blot

Western blotting was performed to assess the time-dependent clearance of *Theileria* parasites. Cell lysates were prepared from cultures after 48 hours of treatment, and protein concentration was quantified. SDS-PAGE and Western blotting were carried out using a rabbit anti-TaSP peptide antibody (1:1000) as the primary antibody, and Anti-IgG HRP-conjugated rabbit secondary antibody (1:2000). β-Actin was used as a loading control (1:5000). Protein bands were visualized using the Bio-Rad ChemidocTM Imaging System with chemiluminescent HRP substrate.

### 2.11 LC/MS-MS and Data Analysis

For mass spectrometry analysis, 100 μg of protein from *T. annulata*-infected cells and cells treated with Altiratinib for 48 hours were extracted. Proteins were reduced with DL-Dithiothreitol, alkylated with Iodoacetamide, and digested with Sequencing Grade Modified Trypsin. The peptides were analyzed using an Orbitrap Fusion mass spectrometer. Proteomic data were analyzed using MaxQuant Software (version 2.6.2.0), with the output files mapped to the proteomes of *Bos taurus* and *T. annulata*. Label-free quantification (LFQ) parameters were used, and the data were processed in Perseus software (version 2.0.9.0). A volcano plot was generated based on LFQ intensity differences, and significant protein hits were selected based on log-transformed values (+/-1.5) and statistical significance (t-test). Best-hit proteins were analyzed for affected pathways using STRINGdb.

### 2.12 Statistics and Reproducibility

Statistical analyses were performed using one-way or two-way ANOVA followed by Dunnett’s multiple comparison test or unpaired t-tests followed by the Mann-Whitney test. GraphPad (Version 9.0.0) was used for all analyses. All experiments were repeated as indicated, and appropriate statistical tests were performed to assess significance.

## 3. Results

### 3.1 Altiratinib binds to conserved PRP4K catalytic motifs in *T. annulata* and other piroplasm parasites for its antiparasitic action

Altiratinib has been shown to inhibit the proliferation of *T. gondii* and *P. falciparum* by selectively targeting the spliceosome kinase TgPRP4K in *T. gondii* and its homolog PfCLK3 in *P. falciparum* [8]. To investigate its potential targets in *Theileria* species of economic significance, we identified two putative uncharacterized protein kinases: TA21325 (serine/threonine protein kinase) in *T. annulata* and TpMuguga_01g00303 (protein kinase domain protein) in *T. parva*, both exhibiting sequence homology with TgPRP4K. Phylogenetic analysis confirmed PRP4K conservation across eukaryotic and protozoan lineages, with distinct evolutionary divergence within apicomplexans. Structural comparisons revealed the conservation of key catalytic motifs, including the adenosine specificity pocket and the DLG motif, across *P. falciparum, T. gondii, T. annulata, T. parva*, and *B. bovis* (Figure_1A, B). Notably, in bovine cells, the DFG motif was conserved, aligning with its human counterpart. The DLG motif, specifically L715 in *T. gondii*, is implicated in Altiratinib binding [8]. Due to the unavailability of a resolved crystal structure for TA21325 (TA-PRP4K), homology modeling was performed using *T. gondii* PRP4K as a structural template. Docking simulations demonstrated that Altiratinib binds favorably to wild-type *T. annulata* PRP4K, yielding a docking score of-13.787 kcal/mol, with hydrogen bonding interactions involving ASP181, LYS64, and TYR117 (Figure_1C).

**Figure_1A.**
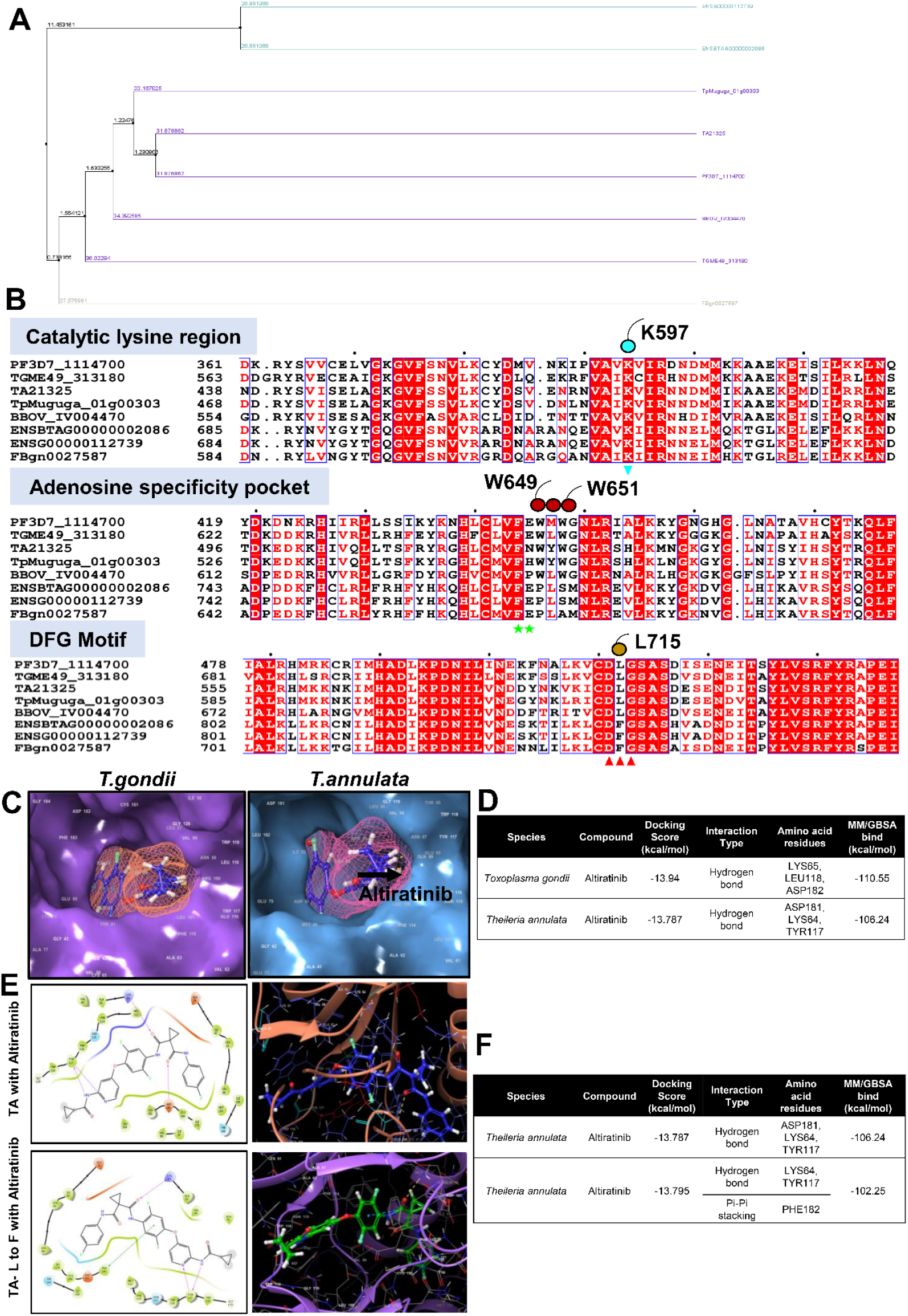
The dendrogram shows that when it comes to PRP4K, the apicomplexans are closer to each other in the phylogeny tree in comparison to other eukaryotes.** Figure_1B**. In the binding of altiratinib, several domains were identified as essential. A comparison with other apicomplexans revealed that the catalytic region and the adenosine specificity pocket are conserved. Additionally, the DLG motif showed similarity across *Plasmodium falciparum (*PF3D7_1114700), *Toxoplasma gondii (*TGME49_313180), *Theileria annulata (*TA21325), *Theileria parva (*TpMuguga_01g00303), *and Babesia bovis(*BBOV_IV004470). In contrast, DFG were conserved across *Bos taurus* (ENSBTAG00000002086), *Homo sapiens (*ENSG00000112739), and *Drosophila melanogaster (*FBgn0027587).**Figure_1C**. Comparison of structure for the altiratinib binding pocket of *Toxoplasma gondii* with *Theileria annulata*. **Figure_1D and 1F**. Docking score and binding affinity values (MM-GBSA ΔG binding score) of both wild type and mutant of *T*.*annulata* protein. **Figure_1E**. *In-silico* images showing wild-type and mutant form (L to F) changes in the binding pocket of altiratinib in *T*.*annulata*.

To assess the impact of structural alterations, a point mutation (L182F) was introduced in *T. annulata* PRP4K. Molecular docking of the mutant protein retained comparable docking scores (-13.795 kcal/mol) but exhibited distinct interaction profiles, including π-π stacking with PHE182 (Figure_1E, F). MM-GBSA-based binding free energy calculations indicated a reduced binding affinity of Altiratinib for the mutant variant (-102.25 kcal/mol) compared to the wild-type protein (-106.24 kcal/mol). Validation studies using *T. gondii* PRP4K (PDB ID: 7Q4A) revealed docking scores and binding affinities consistent with those observed in *T. annulata* (Figure_1D), further supporting the potential of Altiratinib as a PRP4K-targeting inhibitor in *Theileria* spp. Comparison of the conserved sites and the mutations reported in *T*.*gondii* for TgPRP4K (mutations F647S, L686F, and L715F) shows that except for the *B. bovis* parasites, which differ at the L686F position, the rest of the apicomplexan including the *Theileria* parasites, have conserved sites suggesting a possible common mechanism for Altiratinib activity.

### 3.2 *In-vitro* evaluation of Altiratinib against *T. annulata*-infected cells and BPQ-Resistant line

Peripheral blood mononuclear cells (PBMCs) were isolated from *T. annulata*-infected cattle using density gradient centrifugation and subsequently cultured in RPMI-1640 medium supplemented with essential nutrients under controlled conditions. The *T. annulata-*infected lymphocyte cells exhibited continuous proliferation, a hallmark of parasite-driven transformation, validated through PCR amplification of TaSP, a parasite-specific gene.

To assess the therapeutic potential of Altiratinib, it’s *in vitro* efficacy against *T. annulata*-infected cells was examined, with uninfected PBMCs from healthy cattle (confirmed negative for *T. annulata*) serving as controls. Cell viability assays using resazurin demonstrated an IC_50_ value of 6.88 ± 1.5 μM following 48 hours of treatment (Figure_2A). Cytotoxicity analysis in PBMCs from uninfected cattle showed no toxicity at the IC_50_ concentration, and the SI index is also mentioned (Figure_2 B). A time-course assay, measuring viability every 12 hours using trypan blue exclusion, indicated substantial cytotoxic effects on infected cells after 24 hours, with progressive parasite clearance evident in brightfield microscopy over time (Figure_2C, D). Altiratinib was tested against a BPQ-R cell line to further evaluate its efficacy in killing the resistant parasites. Resistance was induced by culturing infected cells in gradually increasing BPQ concentrations, starting at 10 ng/mL. After multiple passages, a resistant strain capable of sustaining growth at 20 μg/mL BPQ was established. This resistance remained stable, persisting even after several weeks in a drug-free environment. Trypan blue exclusion assays demonstrated that Altiratinib effectively blocked the growth of the BPQ-R cells, with brightfield microscopy confirming increased cell death after 48 hours of treatment (Figure_2E, F).

**Figure_2A.**
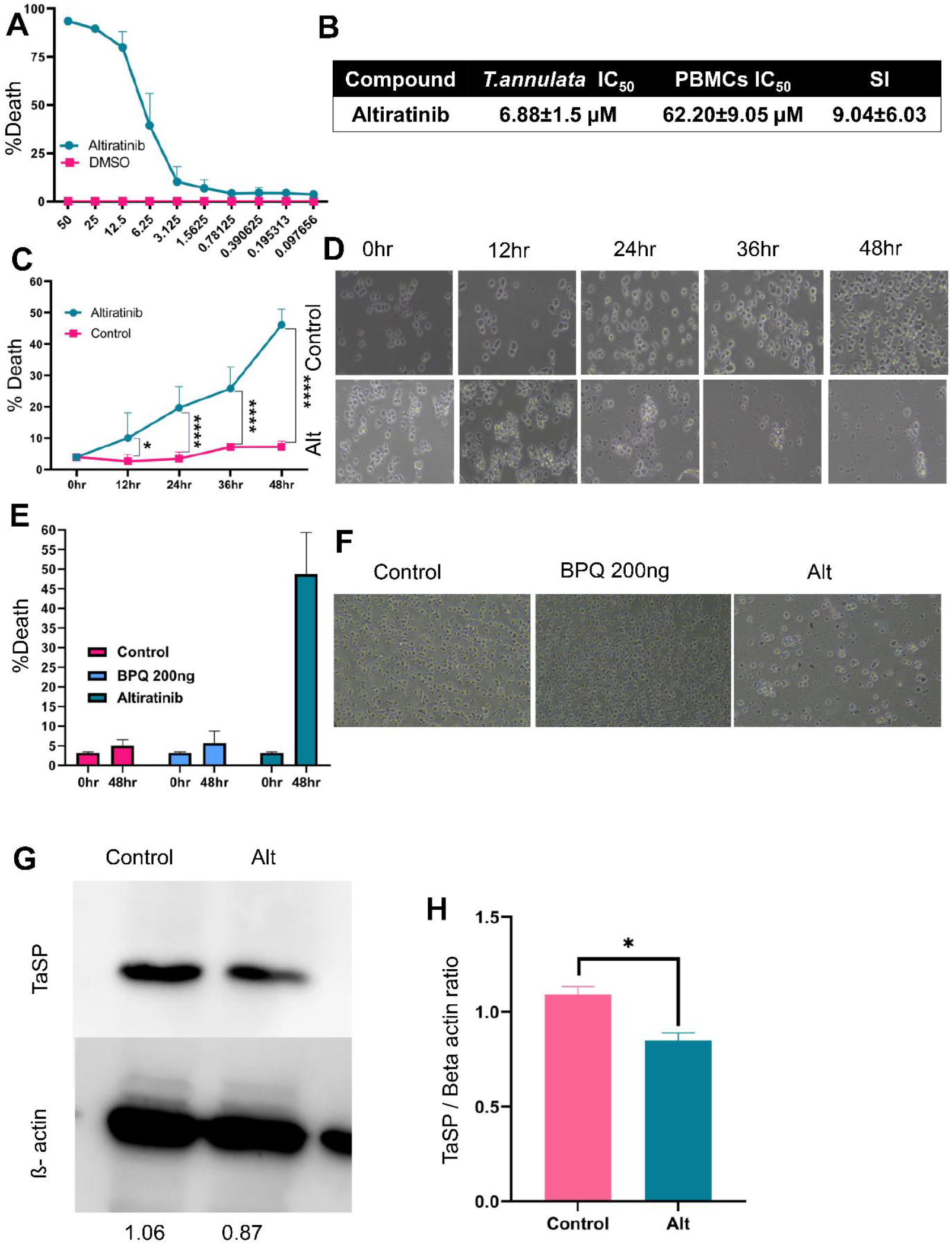
Resazurin assay showing % death in altiratinib treatment *in vitro* compared to DMSO. **Figure_2B**. IC_50_ value of altiratinib in *T. annulata* infected cells and SI index calculated with the help of CC_50_ in PBMCs.**Figure_2C**. A time-kinetics experiment in infected cells demonstrated growth inhibition at 12-hour intervals following altiratinib treatment. In the graph, *represents the significance of (P ≤ 0.05), and ****represents the significance of (P < 0.0001). **Figure_2D**. Bright field images in 20X representing cell morphology changes after treatment. Images are taken at 12-hour intervals time. **Figure_2E**. BPQ-resistant cells treated with altiratinib showing effect post 48hr treatment. BPQ at 200ng is taken as a negative control showing no death in the BPQ-resistant cells. **Figure_2F**. Bright field images of the BPQ-resistant cells. Images taken in 20x represent control, altiratinib at IC50 values, and BPQ 200ng. images are taken after 48hrs of treatment. **Figure_2G**. Analysis of TaSP in *T. annulata* infected cells before and after treatment of altiratinib. Western blotting profile shows a reduction of the TaSP in altiratinib-treated samples. Quantification ratios are mentioned below. ß-actin was used as a loading control. **Figure_2H**. TaSP to ß-actin ratio quantified. In the graph, *represents the significance of (P ≤ 0.05).

Western blot analysis evaluated the specificity of Altiratinib’s antiparasitic activity, targeting the *T. annulata* surface protein TaSP, using β-actin as a loading control. The results demonstrated a significant reduction in TaSP expression following treatment (Figure_2G), with the quantitative analysis presented in (Figure_2H). These findings indicate that Altiratinib effectively inhibits drug-sensitive and BPQ-R *T. annulata* strains *in vitro* while exhibiting no cytotoxic effects on healthy PBMCs.

### 3.3 Altiratinib disrupts host and parasite splicing machinery in *T. annulata*-infected cells

To explore its impact on the splicing machinery in *T. annulata*-infected cells, we conducted LC-MS/MS analysis on both control and Altiratinib-treated cells. Our analysis revealed distinct protein expression changes between control and drug-treated *T. annulata*-infected cells (Figure_ 3A, B). Peptides were analyzed by LC-MS/MS and matched to proteins using species-specific databases for *B. taurus* (UP000009136) and *T. annulata* (UP000001950) in MaxQuant (version 2.6.2.0). The output data was further processed in Perseus (version 2.0.9.0) to remove contaminants and categorical anomalies, retaining only significant peptides in at least all the samples in any group. This analysis identified 6,985 unique host peptides corresponding to 1,220 host proteins. For the *Theileria* parasite, 485 unique peptides were detected, mapping to 99 proteins. The dataset was subsequently used to identify differentially expressed proteins in both host and parasite samples upon drug exposure, applying a log_2_ fold-change cutoff of ±2 and a p-value threshold of 0.01. Treatment led to the downregulation of 64 genes and the upregulation of 2 genes in the host (Figure_3C). Functional annotation identified significant disruptions in biological processes, including mRNA splicing regulation, RNA processing, translation, and ribonucleoprotein complex assembly (Figure_3E). Molecular function analysis revealed altered translation elongation factor activity, RNA binding, and processes related to translational machinery (Figure_3E). Key differentially expressed proteins (DEPs) included ribosomal proteins (e.g., Q76I81, A0A0A0MPA4, Q56JV1, Q2TBQ5, Q3ZBH8, Q58DW5, Q32PB9, P79103), RNA helicases (e.g., F1MPG7, A0A3Q1MR43), and splicing-related proteins such as Q08E38 which is a Pre-mRNA-processing factor 19 (PRPF19) or PRP19. The details are given in Supplementary Table 2.

**Figure_3A.**
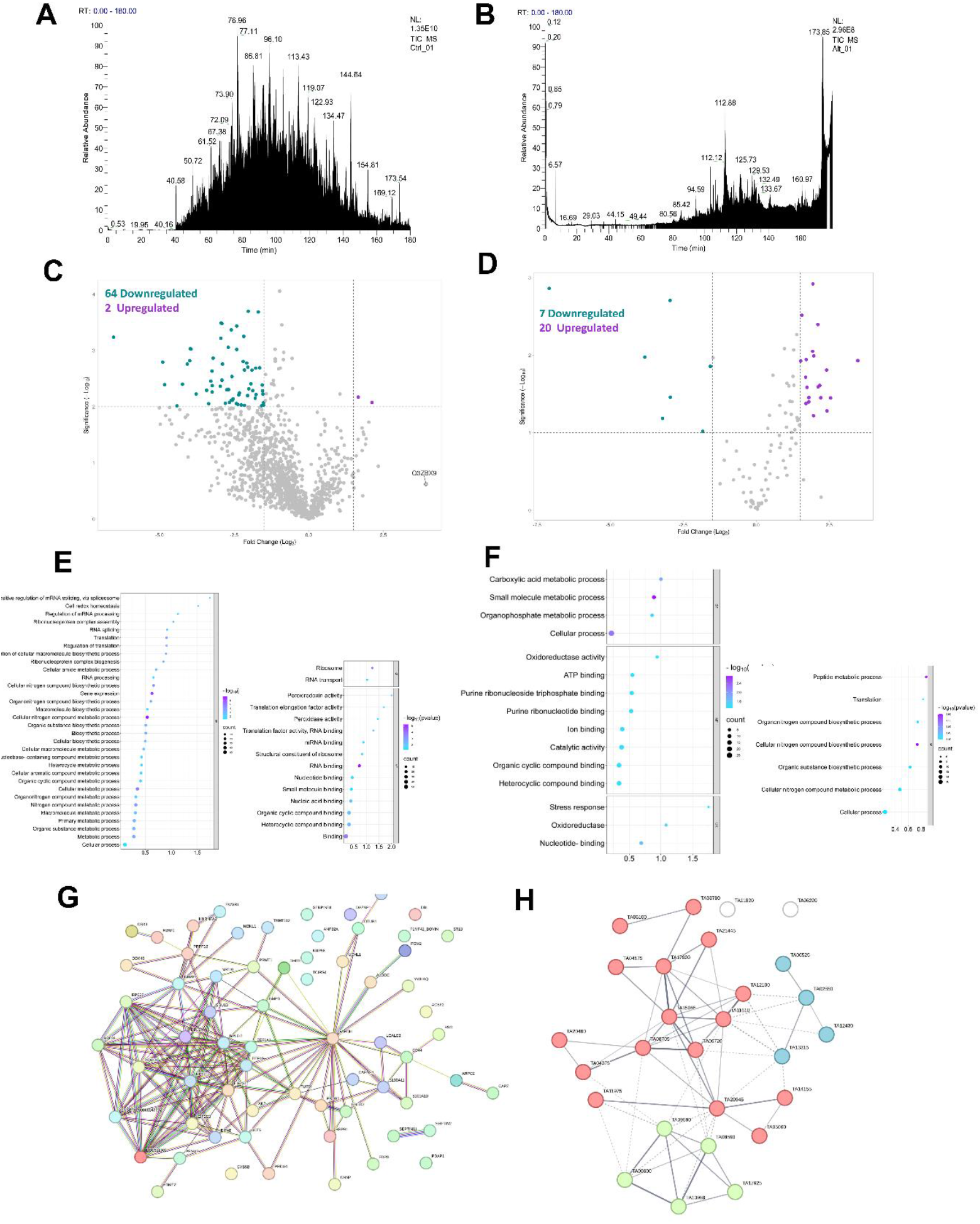
B. To investigate whether altiratinib targets the splicing machinery in the in *Theileria*-infected cells, we conducted a proteomics study, and the chromatograms for both control and treatment are presented. **Figure_3C**. Our study identified 64 downregulated genes and 2 upregulated genes in the host *Bos taurus*. **Figure_3D**. In the parasite *Theileria annulata*, we identified 7 downregulated genes and 20 upregulated genes. **Figure_3E**. In *B. taurus* our analysis of biological function changes revealed differential regulation of processes including positive regulation of mRNA splicing, mRNA processing, ribonucleoprotein complex assembly, RNA splicing, translation, and regulation of translation. In terms of molecular function, we observed alterations in ribosome activity, RNA transport, translation elongation factor activity, peroxidase activity, and various RNA binding activities. **Figure_3F**. In *T. annulata* the analysis identified mechanisms such as carboxylic and metabolic processes, as well as translation-related genes, involved in molecular functions. In biological processes, we observed differential expression of oxidoreductase activity, ATP-binding, stress response, and purine ribonucleotide binding. **Figure_3G, H**. String analysis revealed that most ribosomal proteins are associated with the host, while in the parasite, most proteins are involved in metabolic processes and stress responses.

In the parasite, 7 genes were downregulated and 20 were upregulated following treatment (Figure_3D). Functional analysis linked these DEPs to metabolic processes, translation, oxidoreductase activity, ATP-binding, and stress responses (Figure_3F). Key DEPs included ribosomal proteins (e.g., TA10185, TA14890, TA16000, TA08340, TA08205, TA09180, TA17900, TA09180, TA12815, TA15065, TA21355, TA16165), elongation factors (e.g.,

TA08705, TA05060, TA21190, TA20405, TA06720, TA16305), and splicing-related proteins such as PRP8 homolog (TA03780) and a splicing factor-related protein (TA06100). The details are given in Supplementary Table 3.

Protein interaction analysis revealed that in the host, most DEPs were associated with ribosomal and translational pathways, while in the parasite, they were involved in metabolic processes and stress responses (Figure_3G, H). Altiratinib-induced disruption of translational processes in both host and parasite indicates its antiparasitic efficacy. In the host, key pathways related to RNA splicing, ribonucleoprotein assembly, and translation were significantly affected, likely due to Altiratinib’s targeting of splicing machinery. Notably, splicing-related proteins such as PRP19 in *B. taurus*, PRP8 homolog, and Splicing factor related protein SR protein in *T. annulata* were implicated. The disruption observed in *T. annulata* suggests that Altiratinib directly targets the parasite’s splicing machinery, leading to translational dysregulation. The dual impact on host and parasite splicing pathways supports the hypothesis that Altiratinib exerts its effects by disrupting conserved elements of the splicing machinery, ultimately impairing translational processes in both organisms.

## 4. Altiratinib-induced G1 Phase arrest and inhibited the of proliferation in *T. annulata*-infected Cells

The PRP4 kinase (PRP4K/PRPF4B) family is crucial in regulating pre-mRNA splicing and controlling cell cycle progression across eukaryotic organisms. Following the identification of splicing disruptions in our proteomics analysis, we assessed the effects of Altiratinib on *T. annulata*-infected cells, focusing on cell cycle regulation, proliferation, and signaling pathways.

Treatment with Altiratinib resulted in a significant G1 phase arrest in *T. annulata*-infected cells, as determined by cell cycle analysis (Figure_4A, B), suggesting that the drug impairs cell cycle progression and inhibits the proliferation of infected cells. To further explore the impact on DNA synthesis, we employed the EdU (5-ethynyl-2′-deoxyuridine) proliferation assay, which utilizes copper-catalyzed click chemistry to monitor EdU incorporation into newly synthesized DNA [12]. A marked reduction in EdU incorporation was observed in *T. annulata*-infected cells treated with Altiratinib for 24 hours, indicating decreased cell proliferation (Figure_4C, D).

**Figure_4A.**
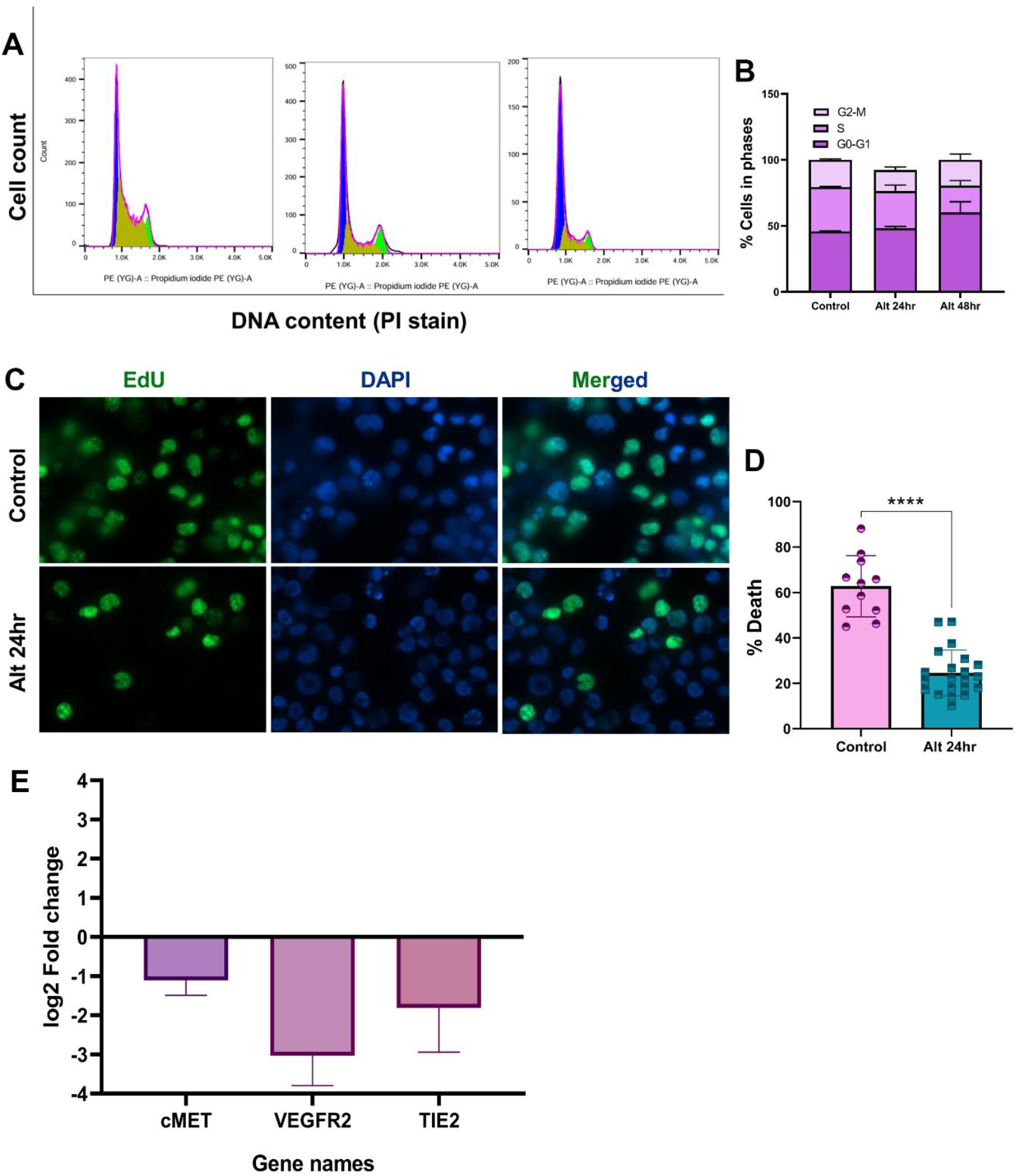
To assess the inhibitory effects of altiratinib on *Theileria*-infected cell proliferation, cell cycle distribution was analyzed 24-and 48-hours post-treatment. PI labeling and flow cytometry showed a time-dependent increase in G1 phase arrest following drug exposure. **Figure_4B**. Cell cycle analysis of both control and treatment representing G0-G1, S, G2-M phase. **Figure_4C**. An EdU proliferation assay performed after 24 hours of altiratinib treatment showed a stall in DNA synthesis compared to the control. **Figure_4D**. Quantification of the Edu proliferation assay. In the graph, ****represents the significance of (*P* < 0.0001). **Figure_4E**. Altiratinib as an anti-cancer drug, inhibits cMET/TIE2/VEGFR2 pathways. To determine if these pathways are affected in *T. annulata*-infected cells, we performed real-time PCR and observed the downregulation of these genes compared to housekeeping genes in treated cells.

Altiratinib, a known inhibitor of the cMET/TIE2/VEGFR2 signaling pathway, has previously demonstrated antiproliferative effects in glioblastoma by targeting this pathway. Given the strong dependence of *T. annulata* parasite proliferation on host cell growth, we examined the effect of Altiratinib treatment on cMET pathway gene expression in infected cells. After treatment, q-PCR analysis revealed downregulation of cMET, TIE2, and VEGFR2. Gene expression levels were normalised to the host housekeeping gene GAPDH (Figure_4E). These findings suggest that the antiproliferative effects of Altiratinib in *T. annulata*-infected cells may be partially mediated through the suppression of cMET signalling.

## 5. Altiratinib triggers ROS-driven DNA damage and mitochondrial dysfunction, leading to apoptosis in *T. annulata*

To explore the molecular mechanisms by which Altiratinib impacts *T. annulata*-infected cells, we assessed its ability to trigger oxidative stress, mitochondrial impairment, and apoptotic pathways. Following Altiratinib treatment, a substantial rise in reactive oxygen species (ROS) generation was observed in the infected cells. The mean fluorescence intensity for ROS in control cells ranged between 68,000 and 70,000, whereas treated cells displayed a significantly higher intensity, ranging from 190,000 to 200,000 (Figure_5A). Increased ROS levels are associated with oxidative damage to essential cellular structures, including DNA [13,14]. DNA damage was confirmed through increased phosphorylation of the γH2AX marker, observed within 24 hours of treatment, indicating significant DNA damage caused by ROS (Figure_5D, E). We further assessed mitochondrial integrity by measuring the mitochondrial membrane potential (Δψm) using JC-1 staining. The red-to-green fluorescence intensity ratio in untreated control cells was approximately 1.5, reflecting normal mitochondrial function. However, after 48 hours of Altiratinib treatment, a significant decline in red fluorescence was observed, reducing the ratio to around 0.5. This shift indicates mitochondrial depolarization and compromised function (Figure_5B, C).

**Figure_5A.**
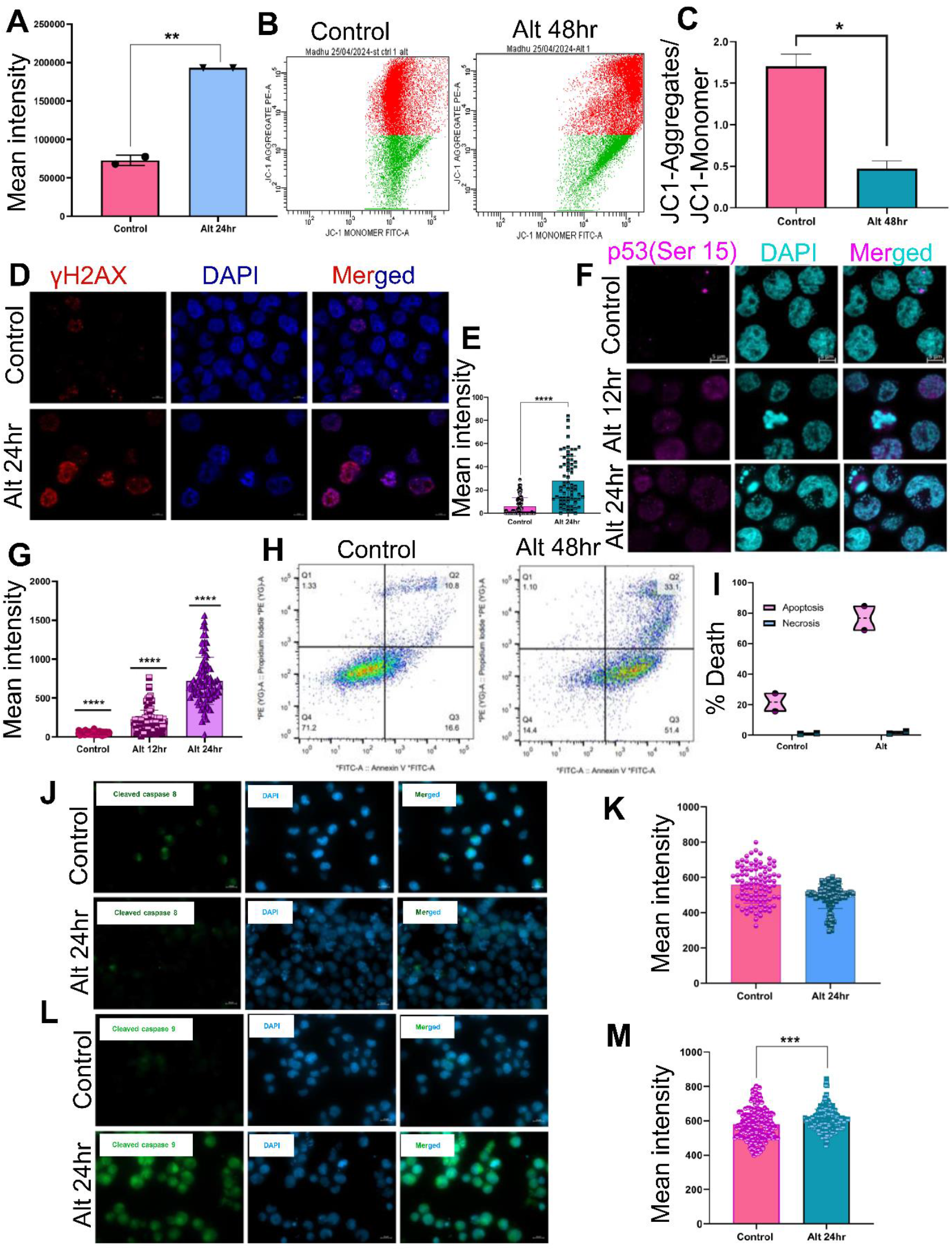
DCFDA assay showing an increase in ROS production in *T. annulata* infected cells after treatment with altiratinib. **Figure_5B**. JC-1 absorption was measured 48 hours post-treatment to assess mitochondrial membrane potential in infected cells. Altiratinib treatment caused a significant reduction in red fluorescence, resulting in a fluorescence intensity ratio of 0.5, indicating mitochondrial depolarization. **Figure_5C**. Quantification of JC-1 assay. **Figure_5D**. We examined the DNA damage marker γH2AX phosphorylated at Ser139 and observed increased levels of phosphorylated γH2AX in altiratinib-treated cells after 24 hours. **Fig_5E**. Quantification graph of phosphorylated γH2AX in both control and altiratinib-treated cells. **Figure_5F**. Phosphorylated Ser-15 P53 being activated in altiratinib-treated *T. annulata*-infected cells. IFA images show there was a significant increase in 12 and 24-hour treatment. **Figure_5G**. A quantification graph comparing phospho-Ser15-p53 levels between control and altiratinib-treated cells. **Figure_5H**. Annexin V-PI assay showing apoptotic cell number increasing in altiratinib-treated samples when compared to control. **Figure_5I**. Quantification showing the increase in apoptosis in altiratinib-treated cells, while cells undergoing necrosis were negligible. **Figure_5J, L**. Altiratinib works through an intrinsic apoptotic pathway. Comparing both cleaved caspase 8 and 9 by immunofluorescence showed there is a significant increase in cleaved caspase 9 after altiratinib treatment. **Figure_5 K, M**. Quantification for cleaved caspase 8 and 9 in both control and treated cells. In the graphs, *represents the significance of (P ≤ 0.05), while **represents the significance of (P ≤ 0.01), ***represents the significance of (P ≤ 0.001), and ****represents the significance of (P < 0.0001).

In response to the ROS-induced damage and mitochondrial dysfunction, we next examined the activation of p53, a critical regulator of the cellular stress response. Immunoblotting for phosphorylated p53 (Ser15) showed activation within 12 hours of Altiratinib treatment, with sustained levels up to 24 hours (Figure_5F, G). To further assess apoptosis, we employed annexin V and propidium iodide (PI) staining, revealing a significant increase in the early apoptotic cell population after 48 hours of treatment, with annexin V-positive cells rising from 7-10% in controls to 68-88% in treated cells (Figure_5H, I). Additionally, immunofluorescence staining revealed activation of caspase 9, a key mediator of the intrinsic apoptotic pathway, and a concurrent decrease in caspase 8 activity, indicating that apoptosis in Altiratinib-treated cells proceeds predominantly via the intrinsic rather than extrinsic pathway (Figure_5J-M).

In conclusion, our findings demonstrate that Altiratinib induces oxidative stress in *T. annulata*-infected cells, leading to ROS-induced DNA damage, mitochondrial dysfunction, and activation of the p53 response. This cascade of events triggers the intrinsic apoptotic pathway, culminating in cell death.

## DISCUSSION

The global rise of drug resistance presents a formidable challenge to human and animal health, particularly within agricultural systems where piroplasm parasites such as *Theileria* and *Babesia* exert a profound economic burden [15]. Tropical theileriosis, caused by *T. annulata*, and bovine babesiosis, attributable to *Babesia bovis*, are major constraints to livestock productivity in endemic regions. Resistance to BPQ—a frontline therapy for theileriosis— underscores the pressing need for innovative treatment strategies that transcend species-specific limitations [16]. In this context, Altiratinib emerges as a promising therapeutic candidate, demonstrating potent PRP4K-targeted activity in *T. annulata*-infected cells. The results support Altiratinib’s potential as a pan-piroplasmic kinase inhibitor, capable of inducing apoptosis and bypassing conventional drug resistance mechanisms in *Theileria* parasites.

The global escalation of drug resistance poses a significant threat to both human health and agricultural productivity [15]. Diseases such as theileriosis, caused by *T. annulata*, contribute to substantial socio-economic losses. The emergence of resistance to BPQ, the primary treatment for theileriosis, highlights the urgent need for alternative therapeutic strategies [16]. This study investigates Altiratinib as a potential antiparasitic agent for *T. annulata*-infected cells, demonstrating its promise as a pan-apicomplexan therapeutic capable of addressing resistance in livestock diseases.

Altiratinib has shown efficacy against other apicomplexan parasites, including *T. gondii* and *P. falciparum* [8]. Comparative sequence analysis revealed conserved motifs, such as DLG, across apicomplexan species. Key residues linked to drug resistance in *T. gondii* and *P. falciparum* (e.g., F647 and L715) were also conserved in *T. annulata, T. parva*, and *B. bovis*. The consistent Altiratinib binding across piroplasms supports a pan-piroplasmic inhibition model for PRP4K, positioning it as a versatile therapeutic candidate for diseases like tropical theileriosis, babesiosis, and potentially East Coast fever. Notably, *B. bovis* exhibited a unique substitution at L686, which may influence its sensitivity to Altiratinib, highlighting species-specific variations in drug response. However, the conserved drug-binding region in *T. annulata* and *T. parva* suggests a mechanism of action analogous to that observed in *T. gondii*. Proteomics analysis revealed that Altiratinib disrupts RNA splicing and translation in both the bovine host and *T. annulata*. Proteomic profiling identified significant alterations in pathways related to RNA processing, mRNA splicing regulation, translation, and ribonucleoprotein complex assembly. The ubiquitin-protein ligase PRPF19, a key component of the PRP19 complex, exhibited differential expression in the host. PRPF19 facilitates spliceosome assembly and stabilization through Lys-63-linked polyubiquitination of PRPF3, enhancing its interaction with PRPF8. It also links transcription, splicing, and DNA repair via associations with RNA polymerase II and the XAB2 complex [17]. Dysregulation of PRPF19 following Altiratinib treatment suggests compromised spliceosomal integrity, leading to defective RNA maturation and translation. Additionally, altered expression of ribosomal proteins and RNA helicases indicates broader disruptions in ribonucleoprotein function, supporting the hypothesis that Altiratinib inhibits RNA splicing. In *T. annulata*, Altiratinib perturbed pathways associated with metabolism, translation, ATP-binding, and stress responses. Differential expression of elongation factors and ribosomal proteins suggests disruption of core translational processes. Key splicing machinery components were differentially expressed, including the PRP8 homolog (TA03780) and a splicing factor-related protein (TA06100). Given PRP8’s central role in the spliceosome’s catalytic core, its altered expression likely contributes to defective splicing and impaired parasite survival [18]. These findings indicate that Altiratinib targets conserved elements of the splicing and translational machinery, with downstream effects differing between host and parasite, potentially due to species-specific regulatory mechanisms. Disruptions in splicing-related pathways in both organisms support the hypothesis that Altiratinib directly interferes with spliceosomal components, impairing RNA processing and protein synthesis. The dual impact of Altiratinib on host and parasite highlights the potential of splicing factors as therapeutic targets for *T. annulata* infections. Further investigation into the molecular interactions between Altiratinib and spliceosomal components is necessary to elucidate its mechanism of action and therapeutic potential.

Given the cancer-like phenotype of *T. annulata*-infected cells, we explored whether Altiratinib targets pathways similar to those in glioblastoma, where it inhibits MET, TIE2, and VEGFR2 signaling [6]. Altiratinib treatment downregulated these pathways in infected cells, as confirmed by RT-PCR, and induced G1-phase arrest and reduced DNA synthesis, consistent with cMET inhibition in glioblastoma. These findings suggest that Altiratinib may impair parasite-induced oncogenic signaling in host cells, enhancing its therapeutic efficacy. Additionally, LC-MS analysis revealed increased oxidoreductase activity and oxidative stress markers in the parasite, indicative of elevated reactive oxygen species (ROS) production. This stress response extended to host cells, may cause mitochondrial dysfunction, elevated γH2AX expression, and intrinsic apoptosis mediated by p53 activation with elevated caspase 9 activity. These findings provide novel insights into Altiratinib’s disruption of host-parasite interactions, sensitizing infected cells to apoptosis. The efficacy of Altiratinib against BPQ-resistant *T. annulata* strains further underscores its therapeutic potential, offering a viable alternative to existing treatments amid rising drug resistance [19].

Future research should focus on *in vivo* validation of Altiratinib’s safety and efficacy in animal models. Pharmacokinetic and pharmacodynamic analyses are essential to optimize dosing regimens and improve bioavailability. Exploring combination therapies with existing drugs may enhance therapeutic outcomes and mitigate resistance development. Additionally, evaluating Altiratinib’s efficacy against other apicomplexan parasites could establish its role as a broad-spectrum antiparasitic agent. In conclusion, Altiratinib exerts its antiparasitic effects by targeting conserved spliceosomal elements, disrupting RNA processing and translation in both *T. annulata* and bovine host cells. It also modulates oncogenic signaling pathways in infected cells, mirroring its effects in glioblastoma, and induces oxidative stress, leading to mitochondrial dysfunction and intrinsic apoptosis. Its efficacy against BPQ-resistant *T. annulata* highlights its potential as an alternative therapeutic strategy. This data validates PRP4K as a druggable and conserved therapeutic node within the parasite kinome, circumventing the need for species-specific drug tailoring. Further studies are warranted to elucidate its precise molecular interactions with spliceosomal components and evaluate its potential in overcoming drug resistance in antiparasitic therapy.

## Declarations

### Ethical Approval

Not applicable.

### Sequence Information

Mass spectrometry proteomics data are available through the MassIVE Consortium using the MassIVE Dataset Submission (1.3.2) workflow with the dataset identifier MSV000097011.

## Author contributions

Most of the experiments were carried out by MS, with support from AS, SS, VMA. *In silico* analysis was done by Si and VB. PS and MS came up with the idea for the study, planned the experiments, and evaluated and interpreted the results. The manuscript was written by PS, MS, Si, AS, and VB. All authors read and approved the final manuscript.

## Acknowledgements

The authors would like to thank the Director (NIAB) for providing all the necessary resources and her consistent support and encouragement. We thank the CSIR JRF grant (AS and SS) and UGC JRF (MS, VMA) for providing fellowship to the PhD students. MS expresses gratitude to the Regional Centre for Biotechnology (RCB) in Faridabad, India, for allowing her to pursue her PhD.

## RESULT 1

## RESULT 2

## RESULT 3

## RESULT 4

## RESULT 5

